# Similarity metric learning on perturbational datasets improves functional identification of perturbations

**DOI:** 10.1101/2023.06.09.544397

**Authors:** Ian Smith, Petr Smirnov, Benjamin Haibe-Kains

## Abstract

Analysis of high-throughput perturbational datasets, including the Next Generation Connectivity Map (L1000) and the Cell Painting projects, uses similarity metrics to identify perturbations or disease states that induce similar changes in the biological feature space. Similarities among perturbations are then used to identify drug mechanisms of action, to nominate therapeutics for a particular disease, and to construct biological networks among perturbations and genes. Standard similarity metrics include correlations, cosine distance and gene set enrichment methods, but these methods operate on the measured features without refinement by transforming the measurement space. We introduce Perturbational Metric Learning (PeML), a weakly supervised similarity metric learning method to learn a data-driven similarity function that maximizes discrimination of replicate signatures by transforming the biological measurements into an intrinsic, dataset-specific basis. The learned similarity functions show substantial improvement for recovering known biological relationships, like mechanism of action identification. In addition to capturing a more meaningful notion of similarity, data in the transformed basis can be used for other analysis tasks, such as classification and clustering. Similarity metric learning is a powerful tool for the analysis of large biological datasets.

## Introduction

Characterizing the biological effects, mechanisms of action (MoA), and phenotypic consequences of drugs is a central challenge in therapeutic development. Relatedly, uncovering the context-specific impact of gene silencing, deletion, or upregulation is a major obstacle to functional characterization in genetics. Biological perturbational datasets, such as the Next Generation Connectivity Map and the Cell Painting project, are tools for these problems that measure the changes in biological features in cancer cell lines due to treatment with small molecules, genetic reagents, and pharmacological agents [1–7]. These data resources can be applied to various tasks like identifying perturbations with similar or dissimilar effects to characterize drug MoAs, identifying candidate inhibitors or agonists of gene targets, and matching therapeutics to disease states [8–10]. These applications require computing similarities between signatures in the given feature space.

The choice of similarity function for biological signatures is often domain-specific, and all are predicated on the assumption that two sets of differential features represent the same biological change if they are sufficiently similar. In gene expression, Gene Set Enrichment Analysis (GSEA), based on the Kolmogorov-Smirnov test statistic, ushered in a new generation of gene set-based similarity functions, including GSA [11], GSVA [12], ssGSEA [13], and others [14–16]. Pearson and Spearman correlation have been used for data modalities such as transcriptomics, phosphoproteomics, and cellular morphology [2, 3, 17, 18]. Cosine similarity, which is mathematically similar to Pearson’s correlation on differentially modulated features, has also been employed to estimate similarities between perturbational signatures [19].

Analysis of perturbational datasets is challenging because phenotypes of interest, e.g., morphology and transcriptomics, have a low signal-to-noise ratio of the measurement assays due to the complexity of the biological systems [2, 4, 20]. Perturbational datasets are commonly used as screening tools to identify candidate reagents for particular tasks, and data generation can be costly. Many analyses use standard off-the-shelf similarity metrics rather than embedding the data in a different basis both because the measured basis has physical interpretation, like transcript abundance or cell shape, and because a standard basis is useful for integration with other biological experiments. Recent studies [17, 21] have found transcriptomic signals that contribute substantially to the variability of the data, can confound analyses, and when removed can improve prediction tasks. This suggests that embedding the data with a data-driven methodology will increase power to uncover biological relationships.

Xing et al formulated a parameterized Mahalanobis distance as a convex optimization problem, bringing the concept of similarity metric learning to the attention of the machine learning community[22]. It has since grown dramatically as a field with a range of applications from information retrieval to bioinformatics[23–29]. A limitation with supervised methods for learning an embedding is that labels are often unknown, difficult to obtain, or intractably costly. Weakly supervised learning (WSL) is a broad term for learning problems where the labels are incomplete or imperfect, and recently it has become a popular method for learning representations on biological data [30–33]. We consider the specific case of WSL of an unsupervised representation learning problem based on replicate labels: two data points belong to the same class if they are replicates of the same experiment, but other associations are unknown. The goal is to learn a similarity function such that similar instances are close while dissimilar instances are far apart, and then do unsupervised learning by applying it to data with unknown class structure. Metric learning with weak supervision relies on replicate data similarly to how contrastive self-supervised learning in the computer vision community relies on synthetic operations that preserve the identity of the samples[34]. Because replicates have to be explicitly measured, the amount of training data available to WSL metric learning is much lower than in the purely self-supervised case. Nevertheless, leveraging replicate data to learn a embedding or new similarity function is a means to learn the inherent properties of the feature space and improve upon static similarity functions.

In this work, we present perturbational metric learning (PeML), a new method for similarity metric learning on biological perturbational datasets that uses biological replicates of experiments as ground truth of similar signatures to learn a similarity function between samples. In contrast with other works, our method is a feature transformation, applied directly to post-processed transcriptomic or morphological features rather than de novo feature extraction from raw data and can be readily applied to different data modalities. We evaluate this similarity function on the L1000 Connectivity Map [4] and the Cell Painting morphological datasets [35, 36]. We show that, compared to conventional metrics like correlation and GSEA-based methods, similarity metric learning (1) improves replicate recall in biological data, (2) improves prediction of compound mechanism of action from perturbational signatures, and (3) can be learned with moderate dataset size, where the number of sets of compound replicates is of the same order of magnitude as the number of features. This method substantially improves the utility of perturbational datasets to extract meaningful biological associations, which will result in better insights into functional characterization of molecules and drug discovery.

## Results

The purpose of similarity metric learning is to find a similarity metric with the best performance at identifying states with biologically similar effects.

To assess the efficacy of metric learning for biological tasks, we applied it to two large-scale perturbational datasets: the LINCS L1000 Connectivity Map[4] and the Center-Driven Research Project (CDRP) dataset of the Cell Painting morphological profiling assay [35]. Similarity metric learning requires a quantitative biological dataset with ground truth that defines elements which should be similar to each other, such as biological or technical replicates. Both the L1000 and CDRP datasets are high dimensional experiments with a rich replicate structure (Figure 1). We compare metric learning to cosine similarity as a baseline using two distinct benchmarks. First, we apply the two metrics to the problem of treatment identification: discriminating replicate signatures of the same compound from signatures of different compounds - the most definitive ground truth in perturbational datasets. Second, we identify compounds with the same mechanism of action - a more difficult problem with weaker ground truth, based on the literature and prior annotation of known, but incomplete, drug mechanisms of actions.

**Fig. 1:**
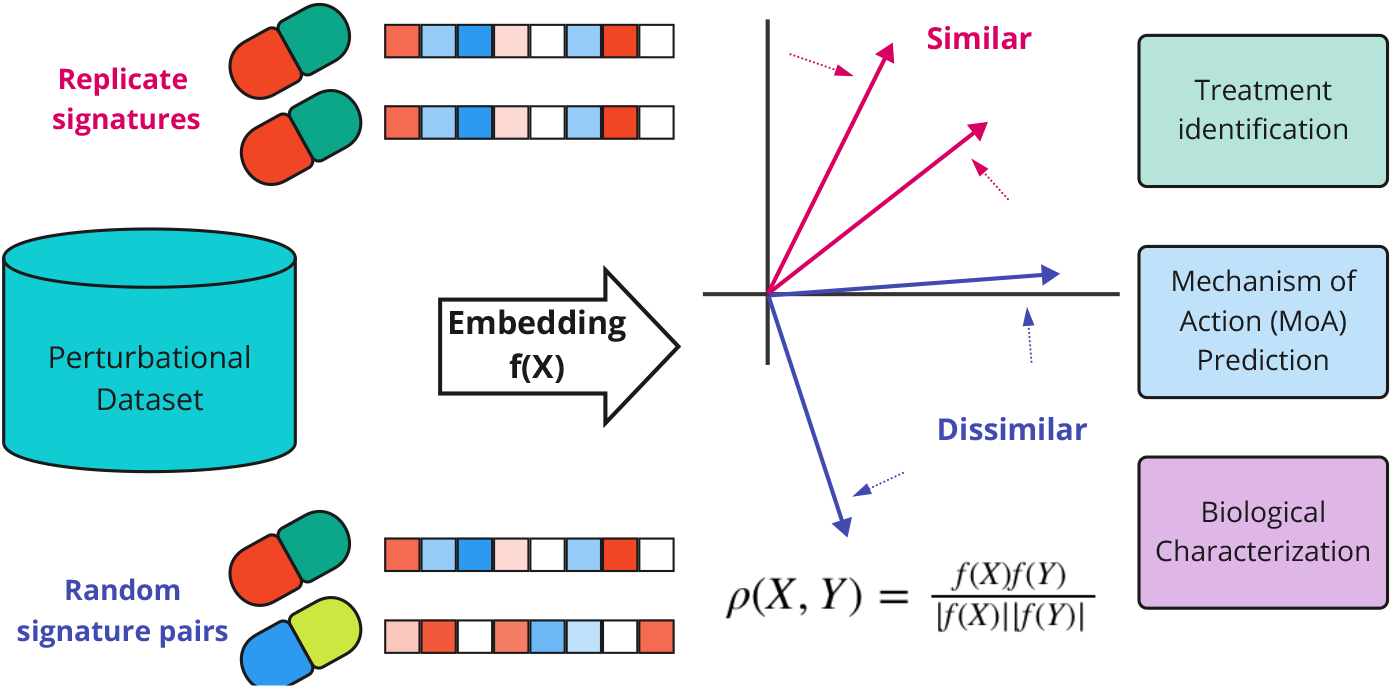
Perturbational Metric Learning (PeML) uses weakly supervised learning to learn a new similarity function on perturbational datasets. The method takes replicate experiments, which are assumed to be similar, and learns an embedding that maximizes the similarity of replicates while keeping random non-replicate signatures dissimilar. The final similarity metric *f* is cosine similarity on the embedded signatures.

### Benchmark Implementation

Both benchmarks are assessed by comparing similarities of positive pairs against the background of the entire population of pairs. For the replicate benchmark, positive pairs are signatures of the same compound treatment, while for the MoA benchmark, positive pairs are signatures of compounds belonging to the same pharmacological class. The problem is challenging because both the positive classes and background (or negative) classes are confounded by elements from the opposite class. While replicate labels for the first task are the gold standard for ground truth signature similarity, the compounds may not be bioactive, may not be active in a particular cellular context, or may be assayed at a time point or dose that does not produce reproducible changes. The ground truth labels for mechanism of action have the same limitations as replicates and may be further confounded by off-target effects. The negative class - the distribution of similarities of random pairs from the population - will contain some pairs which are similar to each other. Nevertheless, a similarity metric that better discriminates positive classes from the background is more useful for detecting biological relationships.

### Performance Evaluation

We evaluate the performance of our metric learning approach using up to three metrics for each benchmark:

1. Average rank (AUC): the rank of positive pairs relative to the background. This is equivalent to the area under the ROC curve or rank curve.
2. Recall at FDR = *α*: Recall (fraction of positive pairs) at a particular FDR
3. Signal-to-noise ratio (SNR) or standardized mean difference: the difference in mean similarity of a positive class and of the negative background normalized by the standard deviation of background similarity

The average rank gives a summary of the overall performance of the similarity metric over all positive classes by computing for each positive pair the fraction of negative pairs with a lower similarity score, and then averaging. While average rank is useful, for practical purposes, we also consider the fraction of positive pairs at a particular false discovery rate. The rank relative to the background can be treated as a p-value, as under the null hypothesis of no association, the rank relative to background is expected to be uniform on [0,1]. By then adjusting for multiple hypotheses with Storey’s False Discovery Rate (FDR) [37], the fraction of positive pairs at FDR *≤* 0.05 is a measure of the power of the similarity metric to discriminate the known positive cases from the background.

However, not all compounds were assayed the same number of times, with the consequence that a small number of compounds constitute the vast majority of replicate pairs (Figure S2). Consequently, we employ balanced sampling to get a more representative measure of performance across classes (see Methods). Balanced sampling allows classes measured frequently to contribute more to the metric while still incorporating information from less measured classes.

### Benchmark 1: Treatment identification and replicate similarity

#### L1000

The L1000 dataset contains differential gene expression signatures of compounds in 230 distinct cancer and immortalized cell lines from a range of tissues, though most of the data are in a few cell lines. Because the underlying biology of each cell line varies, we trained a context-specific model on replicate signatures in each cell line separately. The model is a linear embedding on the space defined by a full rank, positive semi-definite matrix. We train the model with mini-batch stochastic gradient descent, where with each mini-batch, the similarities of a set of replicates are compared to the similarities of a set of non-replicates, and the embedding is updated to minimize the T-loss between the groups (see Methods). To assess whether the similarity metric yields a generalizable representation, we performed 5-fold compound-wise cross validation, so all the signatures corresponding to a particular compound were in either the training set or the validation set.

Metric learning transforms the distribution of similarities of replicates and non-replicates compared to cosine similarity (Figure 2a). The balanced AUC (bAUC) is necessary for measuring performance across a set of compounds which have different numbers of replicates: for those compounds with more than 100 replicate pairs, 100 pairs were randomly sampled, reducing the proportion of replicate pairs coming from any one compound (see Methods). Within each fold of cross validation, the similarities of replicates were ranked against non-replicates, with the expectation that replicates should rank relatively highly, and the overall performance was summarized with an area under the balanced rank cumulative distribution (bAUC). A perfect metric would completely discriminate replicates from non-replicates - with a bAUC of 1 and a balanced recall of 1, while an uninformative metric would have a bAUC of 0.5 and a balanced recall of 0.

**Fig. 2:**
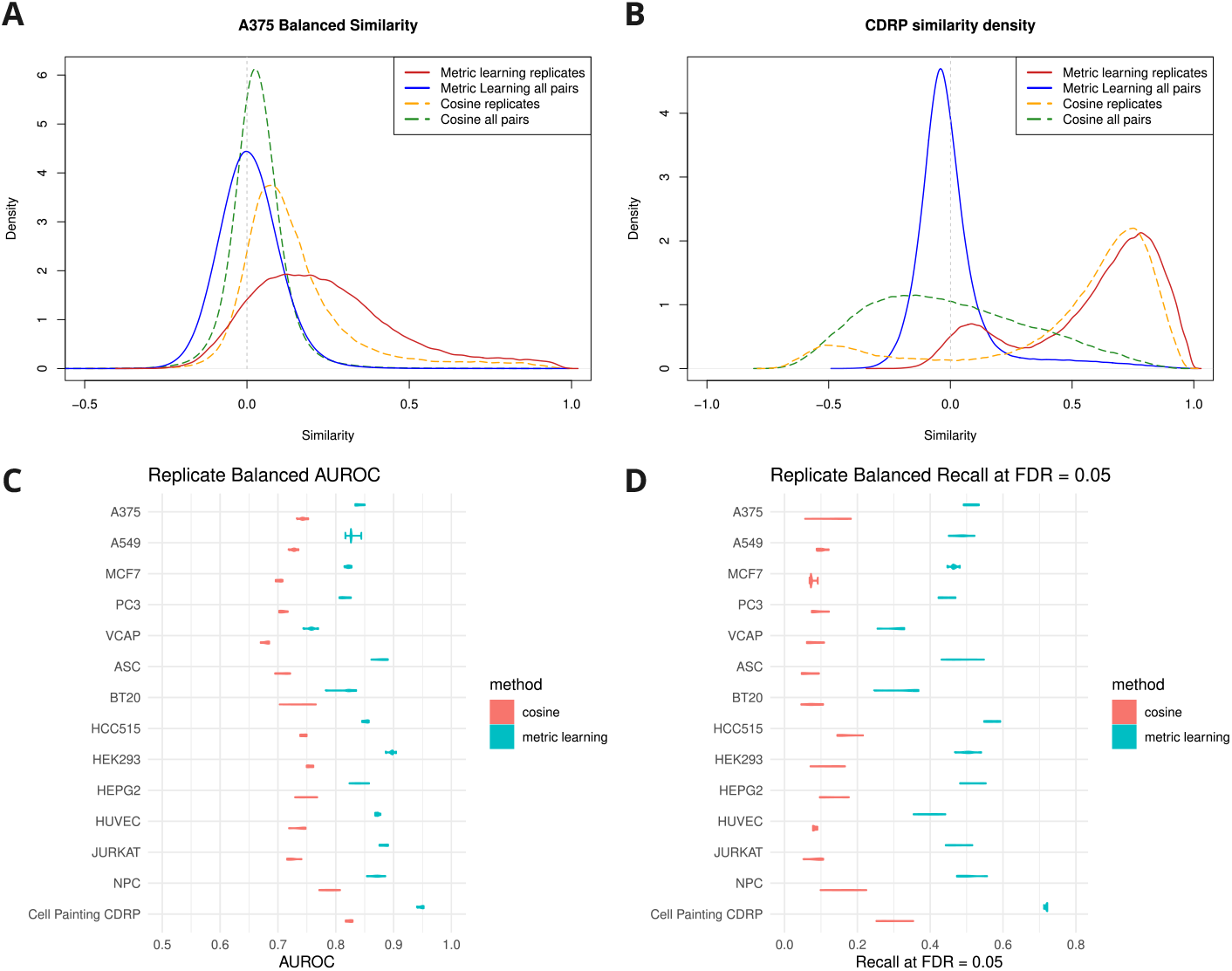
Metric learning evaluation on replicates. (a) In L1000, the distribution of similarities both between replicates and between unrelated compound signatures changes dramatically with metric learning, shown for pairs of replicates and all pairs (non-replicates) in the A375 cell line. Both cosine and metric learning discriminate replicate pairs from non-replicates, but the difference with metric learning is more pronounced. (b) The similarity distributions for Cell Painting replicate data, contrasting cosine with metric learning. (c) Metric learning has improved replicate AUROC over cosine on all cell lines in L1000 and the Cell Painting dataset, measured with cross validation on compounds not seen in training. (d) Metric learning also has improved replicate recall at FDR *<* 0.05.

Metric learning had considerably higher replicate bAUC compared to baseline cosine on test data, with an average bAUC of 0.844 vs 0.732 for metric learning vs cosine, respectively - a substantial improvement in replicate rank (Figure 2c). In addition to average rank, we evaluated the balanced recall of replicates, and found that metric learning recovers a substantially larger fraction of replicate pairs at FDR *<* 0.05 (Figure 2d) in all cell lines tested - with an average balanced proportion across cell lines of 0.462 for metric learning compared to 0.105 for cosine.

### Cell Painting

We then applied the same analysis to the CDRP Cell Painting dataset. Unlike in L1000, the Cell Painting dataset is measured exclusively in U2OS cells, allowing us to include the entire dataset at each level of analysis. Similar to the L1000 replicate analysis, we performed a compound-based cross-validation on all compounds to assess generalizability to unseen compounds. For the replicate recall benchmark, we considered signatures to be part of the same class if they were the same compound treatment irrespective of dose and time point. We find that metric learning transforms the distribution of similarities compared to cosine (Figure 2b). In evaluation on unseen compounds, metric learning has a replicate AUC of 0.947, compared to 0.823 for cosine (Figure 2c). Replicate recall at FDR *<* 0.05 for metric learning is 0.718 compared to 0.299 for cosine (Figure 2d). Balanced recall is not necessary on CDRP replicates, as virtually all compounds were assayed similar number of times (Figure S2). These results show that metric learning is learning an embedding that improves recovery of biological signal and generalizes to data and compounds not seen in the training set in all cellular contexts and in both datasets.

### Benchmark 2: Identifying Mechanism of Action

Demonstrating improvement in replicate similarity and treatment identification benchmarks is necessary to show metric learning is learning a generalizable representation. However, a more pertinent application and a better test of utility is identification of drug mechanism of action, and in particular similarity of signatures of drug treatments with the same MoA.

Subramanian et al [4] defined 92 different pharmacological classes (PCLs) of compounds with sizes ranging from 3 to 44 unique compounds, though not every cell line has data for all the compounds. This PCL collection is not a comprehensive set of compounds with similar classes, but is sufficiently large to serve as a benchmark for biological function. Evaluating the MoA benchmark is similar to the replicate benchmark: positive pairs are signatures of two different compounds belonging to the same MoA class, and negative pairs are those of random compound signatures from the population.

### L1000

We apply the context-specific model trained in each cell line on replicate signatures, and as with replicates, we consider average balanced rank (bAUC) and the balanced recall of same-MoA pairs at FDR *<* 0.05. We find that the average bAUC for metric learning is consistently higher than that of cosine in each cell line (Figure 3A), with an average bAUC across cell lines of 0.677 for metric learning and 0.609 for cosine. This is a substantial improvement in the rank of biologically similar signatures. Additionally, we find that metric learning recovers a greater fraction of same-MoA pairs in all cell lines modeled, with an average recall at FDR *<* 0.05 for metric learning of 0.181 and for cosine of 0.045 (Figure 3B).

**Fig. 3:**
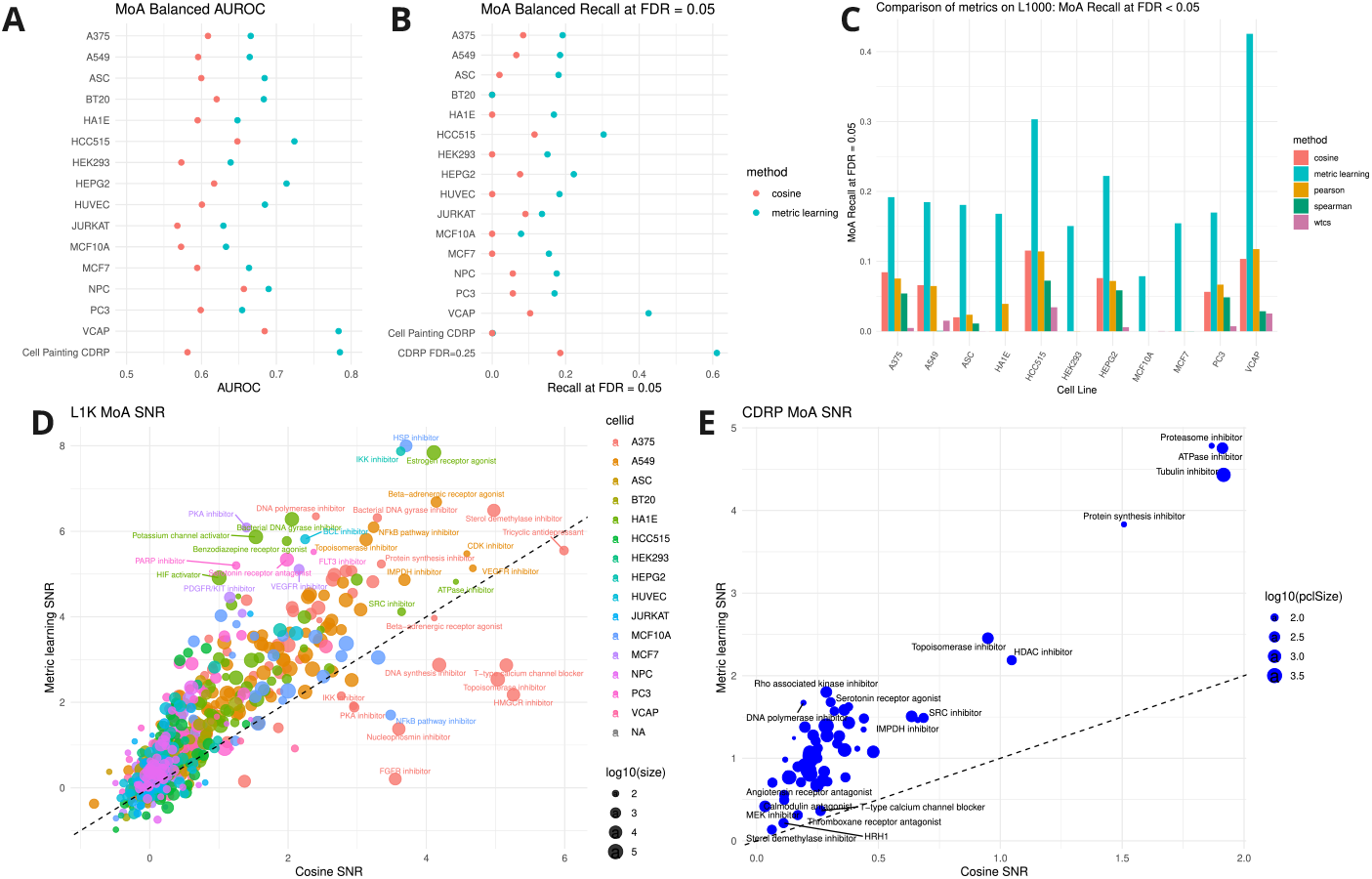
Identifying compounds with the same annotated mechanism of action (MoA). We compute similarities between signatures of compounds with the same MoA and compare relative to the population. (a) Metric learning has improved balanced AUC for signatures of same-MoA signatures compared to cosine in cell lines from L1000 and in Cell Painting data. (b) Metric learning also has superior recall at an FDR *<* 0.05 compared with cosine. (c) In L1000, metric learning has superior MoA recall at FDR ¡ 0.05 than other standard metrics in all cell lines. (d) In L1000, for each MoA x cell line, we compute the signal-to-noise ratio (SNR) with metric learning and with cosine and find that the vast majority of MoAs have improved SNR using metric learning (y-axis) compared to cosine (x-axis). (e) MoAs in the CDRP cell painting dataset, all of which is in U2OS cells, also show improved SNR with metric learning compared to cosine.

While cosine similarity is the most direct comparison for metric learning, we also compared metric learning to other standard metrics used on L1000 including the weighted connectivity score [4], Spearman correlation, and Pearson’s correlation. We find that for the MoA recall task at FDR *<* 0.05, metric learning outperforms the other off-the-shelf metrics in all cell lines tested (Figure 3C).

### Cell Painting

For the cell painting data, as with L1000, we assess the metric performance on biologically relevant mechanisms of action. We use the same groups of compounds as before, and find metric learning has a bAUC of 0.785 compared to 0.581 for cosine (Figure 3A). There is similar improvement in recall at a fixed false discovery rate. At FDR *<* 0.05, metric learning has a recall of 1.46*e* – 3, compared to 2.38*e* – 4 for cosine (Figure 3B). Because so few mechanism of action pairs are recovered in the Cell Painting dataset at FDR *<* 0.05, we also evaluated recall at FDR *<* 0.25 and find metric learning has recall of 0.611 compared to 0.185 for cosine. Importantly, this result shows that metric learning is able to recover a greater proportion of biologically-relevant associations consistently in both the L1000 and Cell Painting datasets.

Both AUC and recall are non-parametric benchmarks, where only the ordering of the positive examples relative to the negative is considered. While ranking is useful, a parametric evaluation - where the values of the similarites of positive pairs are compared to negative pairs - illustrates how well positive groups are discriminated from the background. For cosine similarity and metric learning, we computed the signal-to-noise ratio (SNR) for each mechanism of action relative to the background of all pairs, which is a measure of how well pairs of signatures from that class are discriminated from the population. We found that for the majority of MoA classes, metric learning has greater SNR than cosine in both L1000 and cell painting (Figure 3D, 3E). This parametric evaluation shows that in addition to ranking similar pairs higher than cosine, metric learning better discriminates similar pairs from the background.

### Context specificity

#### Amount of training data

Given the promising performance of metric learning on replicate and MoA recall tasks, we investigated two additional aspects of model generalizability: (1) how much training data is necessary to produce a model that generalizes well, and (2) do context-specific models trained on one cell line perform better on that cell line than models trained in other contexts. To assess how much data is needed, we considered data from four cell lines in L1000, the entirety of the L1000 data, and the CDRP dataset. We excluded 20% of the compounds as the test data with 5-fold cross validation, and trained on the remaining 80%. However, unlike the previous evaluation, we downsampled the training set to ten values between 10 and 3000 compounds, and evaluated the replicate AUC on the test data for each trained model. We compared the test AUC of each model to that of cosine to determine how many sets of training replicates are necessary to improve above cosine. We found that while there is some variability, our metric learning approach yielded better AUC on all the datasets with 300 training compounds, and some improved over cosine with as few as 50 compounds (Figure 4A). Importantly, this suggests that datasets with only a few hundred conditions with replicates can use metric learning to improve biological recall.

**Fig. 4:**
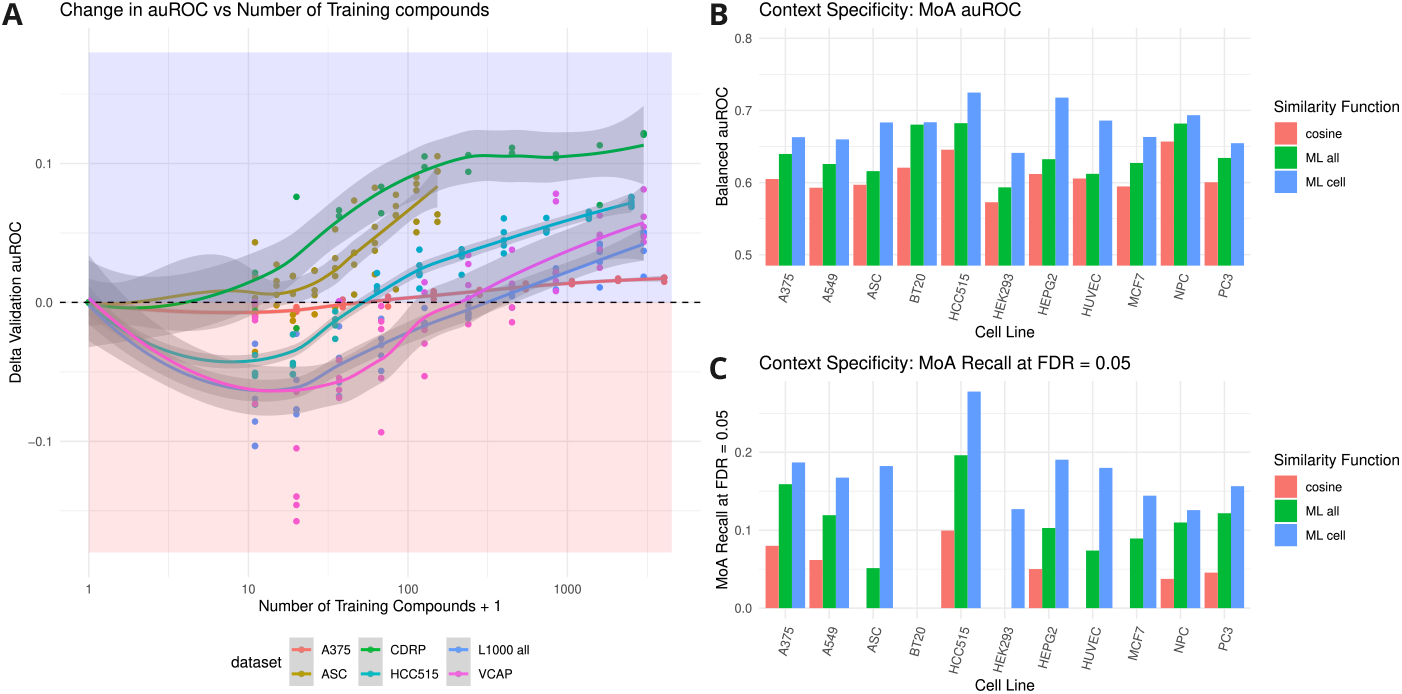
Training data size requirements and context. (a) The model generalizes better to unseen test data with increasing number of training samples (x-axis), as measured by change in auROC relative to cosine (y-axis). The red region indicates the regime where cosine outperforms metric learning, and blue where metric learning outperforms cosine. With small numbers of compound classes (*<* 100), metric learning performs worse at compound query recall than with base cosine (x=0). However, above about 100-500 compounds, the model’s auROC exceeds cosine. (b, c) Metric learning functions learned on cell line-specific data outperform a pan-dataset function on MoA prediction. (b) MoA auROC in a cell line is higher for the cell-line specific function compared to the pan-dataset function, and both metric learning functions outperform cosine. (d, e) MoA Recall at FDR *<* 0.05 on a cell line is greater using a cell line-specific function compared to a pan-dataset function, both of which outperform cosine.

#### Cell context

One of the modeling premises was that metric learning would work best with a different model for each cancer cell line in L1000 due to intrinsic differences in the biology of those cell lines. To evaluate this, we trained a pan-cell line metric learning similarity function on the entire L1000 dataset, where signatures were treated as replicates if they were of the same compound in the same cell line. We then compared cell line-specific metric learning functions to the pan-cell line function on the task of identifying pairs of signatures of compounds with the same MoA. We used cosine as a baseline, and evaluated with balanced AUC and balanced recall with FDR *<* 0.05, as before. We found importantly that while both the pan-cell line metric and the cell-specific metric had better performance than cosine, the cell-specific model showed greater improvement. The mean improvement in MoA AUC over cosine was 0.029 for the pan-cell line metric and 0.070 for the cell-specific metrics (Figure 4B). The mean improvement in recall at FDR *<* 0.05 over cosine was 0.062 for the pan-cell line metric and 0.137 for the cell-specific metrics (Figure 4C). This result suggests that while metric learning may be applied to a dataset with heterogeneous composition effectively, learning a context-specific model on subsets of data with consistent underlying biology can result in further improvement at similarity retrieval tasks.

#### Properties of the Embeddings

To understand how the metric learning embedding is altering the space, we ran PCA on the original data space and on the embedded space for each cell line in L1000 and for the CDRP dataset and considered the distribution of eigenvalues (Figure 5). The distribution of eigenvalues of the principal components (PCs) in the base (unmodified) and embedding (metric learning) space characterizes the underlying dimensions of the spaces. Three key differences about the eigenvalue distributions are immediately apparent.

**Fig. 5:**
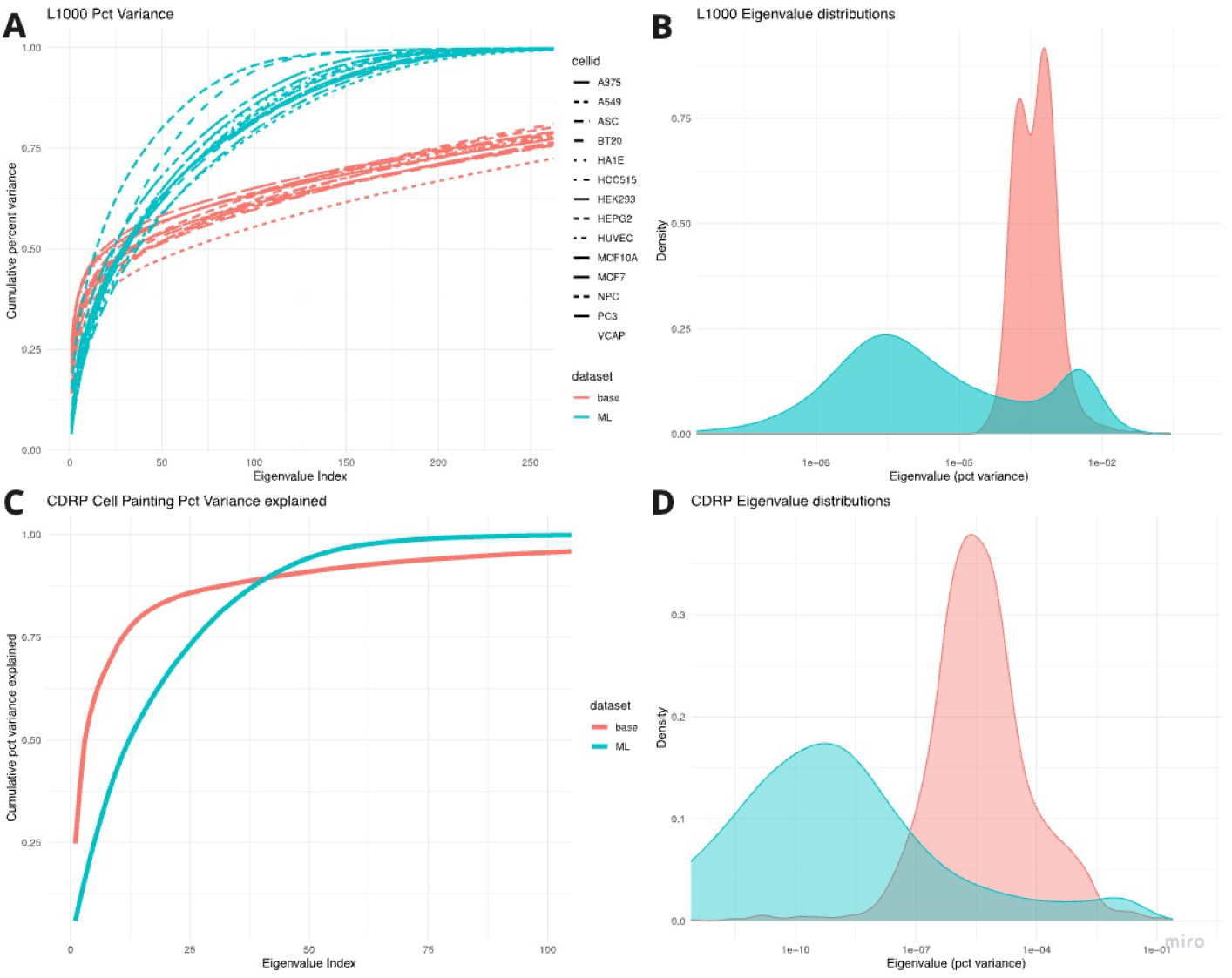
Distributions of the eigenvalues of the base and embedded spaces. (a) Each line corresponds to the cumulative percentage variance explained by the first N principal components for a cell line in the unmodified space (red) and in the PeML embedding (blue). (b) The percent variance explained by each of 978 principal components for all cell lines is displayed as a probability distribution function to highlight differences in the density between the two spaces. (c,d) The cumulative variance explained and probability distribution of percent variance explained for the CDRP cell painting base and metric learning embedded spaces.

First, the embedding uses only the first 200-250 PCs to explain all the variance, whereas the base space requires additional dimensions. On average across cell lines, for L1000, the metric learning embedding space explains 98.6% of the variance with the first 200 PCs, while the base space explains only 71.9%. On the CDRP dataset, the effect is similar though less pronounced, with the first 100 PCs explaining 99.8% of the variance for the embedding and 95.7% for the base space. We surmise that many of the dimensions that are effectively eliminated from the embedding are biological noise signals that do not help discriminate repeated experiments, suggesting that the metric learning embedding helps denoise the respective measurement spaces.

Second, the largest eigenvalues are relatively larger in the base space than in the embedded space. Averaged over the cell lines in L1000, the first five PCs explain 32.8% of the variance in the base space and 20.7% in the embedded space. On the cell painting CDRP data, the first five PCs explain 60.2% of the variance in the base space and 25.3% in the embedded space. This result is consistent with other studies [17, 21] that have shown the presence of large scale confounders, and that reducing the relative contribution of the largest signals improves power.

Third, the non-zero eigenvalues are more equally distributed in the embedded space. We restricted our analysis to the first 200 PCs for L1000 because the embedding reduces subsequent eigenvalues to effectively zero (Figure 5b, d). For L1000, we found that the gini coefficients of the first 200 eigenvalues average 0.559 in the embedded space and 0.640 in the base space. For cell painting, the gini coefficients for the first 100 PCs were 0.652 for the embedded space and 0.819 for the base space. The embedded gini coefficients are lower (p = 1.7*e −* 3 by paired Wilcoxon test) (Figure S5). In summary, this result suggests that the metric learning embedding is more equally distributing the contribution from the informative biological signals while simultaneously denoising the signals.

## Discussion

In this work, we designed and applied PeML, an implementation of weakly supervised learning to perturbational datasets from two different modalities - gene expression and cellular morphology - and demonstrated improved recovery of known biological signals. Importantly, the improvement was observed both in unseen compound replicates and in compounds with similar mechanism of action, which demonstrates that metric learning provides a generalizable similarity metric that more faithfully represents biological similarity. We demonstrated that metric learning improves both the ranking and the power to discriminate similar pairs from the population using parametric and non-parametric benchmarks. The same metric learning methodology was used on both transcriptomic and cell morphology signatures - which are significantly different feature spaces, illustrating the versatility of the method and its robustness to feature variability and covariance.

Weakly supervised learning requires only replicate data for training without additional annotation or prior knowledge, meaning that the method can be applied to many datasets where the only labels are reproduced experiments, eliminating the requirement for additional labels. As the scale and complexity of perturbational biological datasets grows, the capability to learn a representation that improves retrieval and discovery of associations among data points based solely on treatment metadata without further annotation is powerful. Furthermore, because the underlying signal in these datasets is small compared with the noise, tools that can extract more information from the data solely with computational methods dramatically improve the utility of the data. Our method also has the advantage of being applicable to processed datasets rather than requiring generation of features from raw data.

We also demonstrate that context-specific models trained on an individual cell line have superior performance for matching biologically-related signatures than models trained on the entire dataset or on different cell lines. This finding suggests that there are important biological differences between cell contexts that are captured by the embeddings. The result on the amount of data needed to learn a useful embedding can guide whether sufficient data exists for a particular cell line to train a context-specific embedding that can outperform a pan-dataset embedding.

While the subject of this work was metric learning on perturbational datasets, we surmise that the general method of weakly supervised learning can be used on other datasets to learn intrinsic similarity or distance functions for subsequent analysis. For example, analysis of single-cell sequencing data in some feature space could be a promising application for feature-specific intrinsic similarity functions provided a notion of ground truth can be obtained, for instance from complementary assays characterizing some biological phenotype.

The more general self-supervised contrastive learning has sparked a revolution in the machine learning community, especially in computer vision, but it relies upon synthetic, identity-preserve operations or data augmentations to generate pairs of samples with the same class. For imaging, this can be done with rotation, rescaling, jitter, distortion, or censoring [34]. A major obstacle to applying these methods to biological data, especially molecular features, is that the data augmentation operations that preserve the identity of samples are not obvious. The space of transformations on transcriptomic data, for example, that leave the identity of the biological state unchanged is unknown. Using an obvious transformation like Gaussian noise is only a poor approximation of the neighborhood of a sample on a biological manifold. Until effective synthetic data augmentation operations can be demonstrated, replicates will have to suffice as a method to obtain similar pairs.

In summary, metric learning using weakly supervised learning is a powerful tool for maximizing the utility of perturbational experiments that does not require labeling besides experimental metadata. The method learns an intrinsic similarity function specific to the cellular context and nuance of each feature space. The method generalizes to more than one feature space and consistently improves identification and recall of biological relationships.

## Methods

### Perturbational Datasets

The next generation L1000 Connectivity Map was first published in 2017 as part of the NIH Library of Integrated Network-Based Cellular Signatures (LINCS) Consortium [4]. The data consists of differential expression signatures measured as a result of treatment by small molecules and genetic reagents in 978 measured landmark genes and 11350 inferred genes. For our analysis, we used level 5 compound data and only used the 978 landmark genes. The inferred genes are all imputed with a linear model on landmark genes, so the landmark genes contain all the information about the signature. We used the standard dataset with normalization and data processing pipelines as discussed in Subramanian et al. We accessed the Expanded CMap LINCS Resource 2020 (CMap 2020) via the Broad’s web portal at http://clue.io.

The Cell Painting dataset we used was the Center Driven Research Project, first published in 2017 [35]. The dataset consists of 153,370 signatures of 30,412 compounds assayed in the U2OS cancer cell line, and the values are changes induced by the compound in 1783 engineered morphological features extracted with CellProfiler. The data was downloaded from http://gigadb.org/dataset/view/id/100351/File_page/42, and the consensus profiles were used in the analysis.

### Similarity Metric

Cosine similarity on two vectors *A, B ∈ R*^*N*^ is given by:

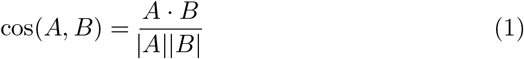

We propose to learn a similarity function *f* from a class of rescaled inner products parametrized by the square matrix M where the similarity between two vectors *A* and *B* is given by:

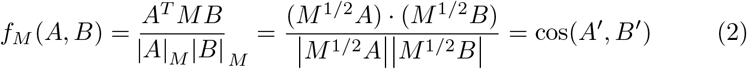

Where 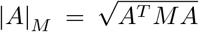, i.e., the Euclidean norm in the embedded space. Equivalently, this is the cosine distance on an embedding given by *A*^*′*^ = *M* ^1*/*2^*A*. The learned metric is then both a natural distance function on the original space and an embedding into a natural basis, i.e. the basis of the dataset that more faithfully recapitulates known relationships. In principle, non-linear embeddings or embeddings changing the number of dimensions can be trained with the same procedure.

To achieve this, we apply weakly supervised learning, which requires pairs of data points that are known to be similar to each other. By assumption, they should be more similar to each other than they are to other data points in the space. Metric learning seeks to maximize similarity between *a priori* similar points while minimizing similarity between dissimilar points. Given a set of similar points *S* and random pairs of elements *E*, we then choose M to minimize the loss function *L*, where *μ*_*E*_ denotes the mean similarity of *f*_*M*_ over pairs of points in *E* (analogously for *μ*_*S*_), and *σ*_*E*_ denotes the standard deviation of *f*_*M*_ over pairs of points in E.

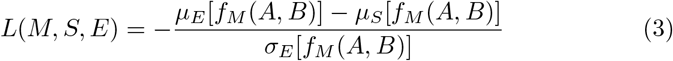

This loss function is related to the t-statistic of the similarities on the set *S* compared to those on the set *E*, hence t-loss. This t-loss is similar to cosine loss [38], but importantly normalizes by the standard deviation of *f*_*M*_ over the space of unrelated pairs. It is equivalent to maximizing the z-scores of similarities of similar pairs. We minimize the loss using mini-batch stochastic gradient descent with the Adam optimizer, implemented in torch for R (torch 0.10.0, available on CRAN). For the perturbational data, each mini-batch consists of a set of similar signatures corresponding to all signatures of one compound treatment, and the set of dissimilar points are a random sample of signatures of twice the size of the similar signatures.

Unless otherwise specified, the entire set of compound treatments for a particular cell line was used for training. Within an epoch, each compound with more than two replicate signatures was seen in training once as a mini-batch. For a mini-batch, the positive class consisted of signatures of a particular compound, and the negative class consisted of twice as many random pairs of signatures; for instance, a mini-batch might consist of ten signatures of a compound compared against twenty random signatures. The positive class was then similarities between the compound replicates, and the negative class was similarities between the random pairs. The adam optimizer used a learning rate of 0.01, which was chosen from cross validation. The final output is an embedding which linearly transforms the underlying data space with a positive semi-definite matrix, and the learned similarity function - cosine similarity on the embedding - follows naturally.

### Benchmarking procedure

In this paper, the metric learning similarity function (or equivalently, embedding) was trained with positive pairs as signatures of the same compound in the same cell line irrespective of dose and time point, and negative pairs as signatures of different compounds in the same cell line.

To evaluate whether metric learning was generalizing to unseen data, we used cross-validation on replicates. Each dataset (L1000 cell lines, the entire CDRP dataset) was divided into five folds by compound, so all the signatures of any compound were in the same fold. The metric learning function was trained on four folds and then evaluated on the excluded fold. Any signatures used in the replicate benchmarks were unseen in the training process and were of compounds that were not used in the training process.

### Benchmarking metrics

We considered three metrics for benchmarking, all of which rely on comparing elements of positive classes to a negative class. As discussed, the classes are imperfect. For replicates, the positive class consisted of pairs of the same compound in the same cell line irrespective of dose and time point, while negative pairs were signatures of different compounds in the same cell line. This is imperfect because there are examples of positive pairs which lack biological similarity - for instance, if the dose is too low to induce phenotypic changes, the time point poorly timed to capture the changes, or the compound not bioactive in that cell line. Conversely, in the negative class, there are examples of pairs of compounds with the same MoA but not annotated as such, with the same off-target, or activating similar broad transcriptional programs (e.g., apoptosis). Nevertheless, these class definitions are the best available notions of ground truth that do not rely upon using the data to assign labels.

For the replicate benchmark, positive classes consist of similarities between replicate signatures, where each class corresponds to a single compound. For the MoA task, positive classes consist of similarities between groups of signatures with the same annotated mechanism of action. We used two non-parametric benchmarks that consider only the ranking of positive class members relative to negative, and one parametric that compares the similarity values.

1. Balanced auROC (bAUC): the area under a balanced ROC curve, or equivalently, the average rank of positive examples relative to negative ones. Because the number of signatures per compound and cell line varies dramatically - from 1 to 813 (for the positive control bortezomib in MCF7 cells), we used a *balanced* ROC curve. For the balanced auROC, for each positive class, we sample whichever is smaller of the number of pairs or a parameter N of pairs, then construct an ROC curve in the usual way. We choose N = 100 for replicate analysis in this work and N = 1000 for MoA classes. For compounds with few examples, all of the pairs are then included. For compounds with more than N pairs, only N pairs are included. With a standard ROC curve, the distribution of signatures per positive class is so skewed that as few as two or three compounds account for more than 50% of positive pairs (Figure S2). With the balanced ROC curve, the importance of each class is more equally weighted. Compounds that are measured a small number of times are given less weight than those measured 15 or more times, reflecting the lesser confidence in few pairs of measurements.
2. Recall at FDR = *α*: the fraction of positive pairs with a q-value *< α*. As before, we used balanced sampling so that the benchmarking metric is not dominated by a small number of positive classes. First, positive pairs are ranked relative to the distribution. As under the null hypothesis of no association, this ranking would be uniform, the rank can be treated as a p-value. We then balanced sample the p-values, so positive classes with more than 100 pairs are downsampled to 100 examples. Finally, we compute a multiple hypothesis-corrected p-value using the FDR procedure, and assess how many of the positive pairs have a q-value below the threshold *α*. Practically, this indicates what fraction of positive pairs would be identified as associated if the positive examples were interrogated for similarity. This does not represent how many of the positive pairs would be discovered if they were not explicitly considered, i.e. if a similarity matrix were computed among all pairs of signatures and the top K most similar pairs identified as associated.
3. Signal-to-noise ratio (SNR) or standardized mean difference: 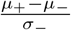. For each positive class, compute the mean intragroup similarity: 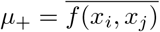, where *x*_*i*_, *x*_*j*_, *i* ≠ *j* are elements of the group, and f is the similarity metric of interest. This is then standardized by subtracting the mean negative group similarity *μ*_*−*_ and standard deviation of the similarity of the negative group, *σ*_*−*_. For replicates, the negative group consists of pairs of signatures of different compounds; for the MoA task, the negative group is pairs of signatures of compounds not belonging to the same MoA. The SNR gives a parametric evaluation of how different the intragroup similarity is for each group compared to the background, essentially a z-statistic of the intragroup similarity. Larger values indicate that the group is better discriminated from the background, and 0 indicates no discrimination.

### Benchmark 1: Replicate similarity

For the replicate similarity benchmark, because the metric is learned by forcing replicates to be more similar to each other, we used five-fold compound-wise cross validation. For each fold, the signatures of one fifth of compounds were held out for validation, and the remaining 80% were used for training in the usual way. We then evaluated the benchmark metrics on the validation fold. This is analogous to applying a PeML similarity function learned on a dataset to signatures of new compounds drawn from the same distribution of compounds. To assess improvement, we compared PeML to cosine, because cosine is the base case of PeML with an identity embedding.

### Benchmark 2: Identifying Mechanism of Action

The mechanism-of-action benchmark applies the similarity metric to pairs of signatures from different compounds but belonging to the same pharmacological class, and so none of the MoA positive examples are ever used as positive pairs in training of the similarity metric. We therefore first trained a similarity metric using the entire dataset of replicates, and then applied it to the MoA task. We used the Pharmacological class mappings from [4], which were obtained from the literature and public databases. This is not a comprehensive list of annotations, but was used for consistency and to provide a thorough assessment of similarity metric performance. As with replicates, the different MoA classes had different sizes. We used balanced sampling with N=1000 to account for the larger sizes of MoA classes and compared with cosine. As with replicates, we compared same-MoA similarities against the background of all pairwise similarities in the same cell line to generate ranks for benchmarks and to compute the signal-to-noise ratio.

### Research Reproducibility

The data is publicly available at the locations listed above. The code used for the analysis in this paper is available in https://github.com/bhklab/metriclearning, and the PeML method will be made available as an R package.

## Glossary of Terms

bAUC: balanced area under Receiver Operating Characteristic curve (auROC)
CDRP: Center-Driven Research Project, the name of a Cell Painting morphological dataset[35]
L1000: Luminex L1000, the name of an assay used for the Connectivity Map, or equivalently the dataset generated by that assay
MoA: mechanism of action
PCA: Principal Component Analysis
PeML: Perturbational metric learning, the method introduced by this paper
WSL: Weakly supervised learning

## Supplemental Figures

**Fig. S1:**
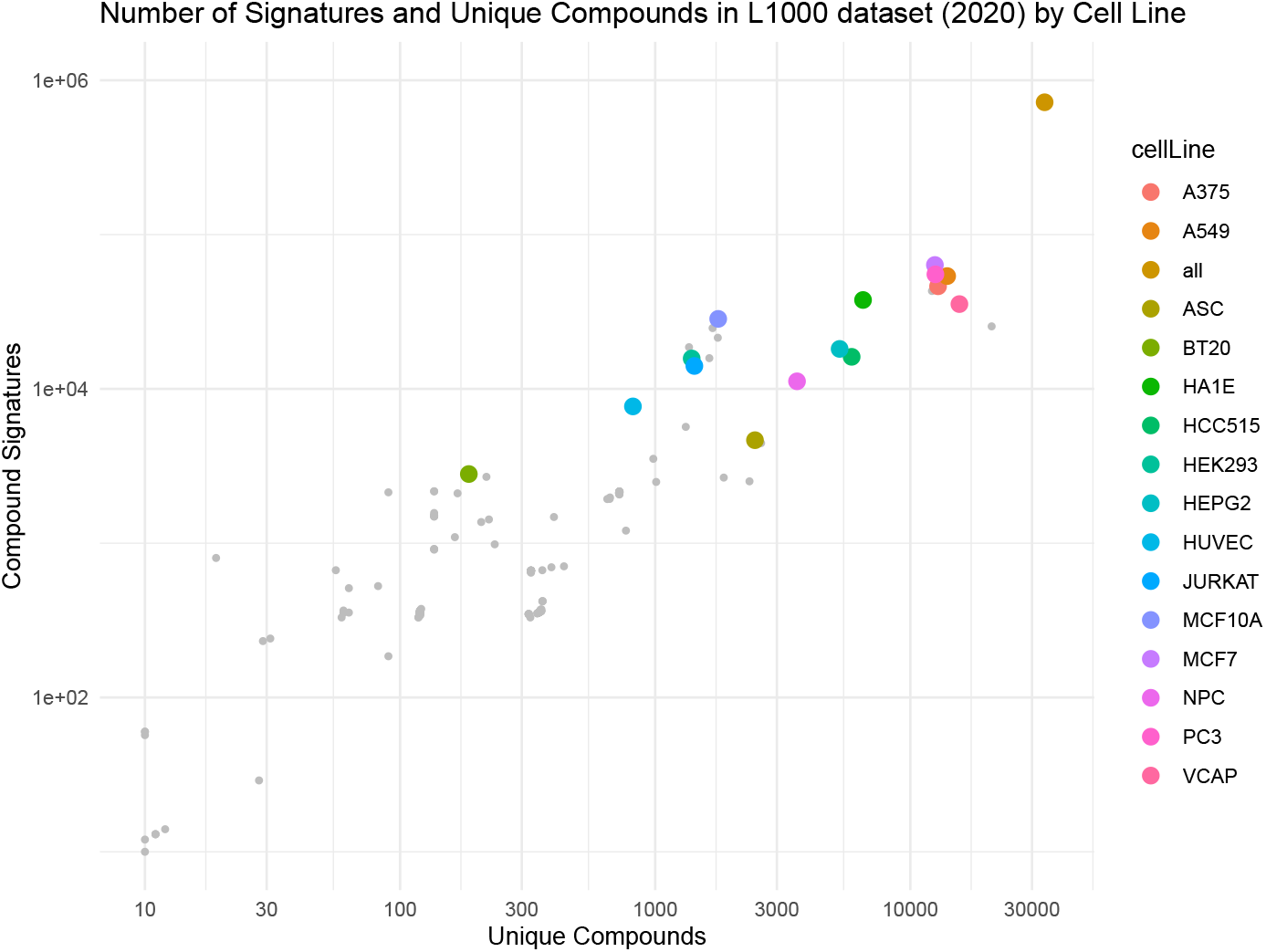
Count of signatures and unique compounds in the cell lines in the L1000 dataset used in this analysis. The colored points are the cell lines used in this analysis, and the grey dots correspond to other cell lines that were not modeled.

**Fig. S2:**
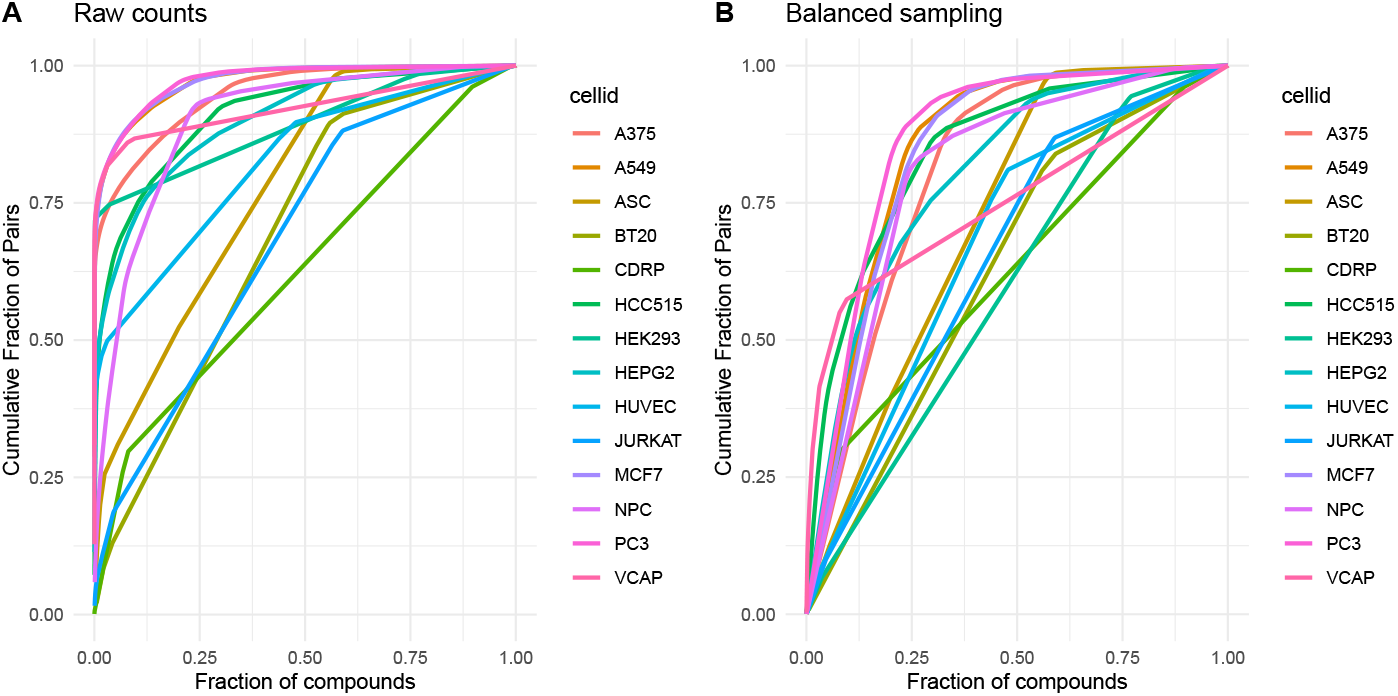
Balanced sampling is necessary for L1000 replicates to mitigate the non-uniform distribution of compound replicate counts. Individual compounds are assayed different numbers of times in the L1000 dataset, and the number of pairs of replicate treatments for a compound goes as *C*(*N*, 2) *∼ O*(*N* ^2^). Thus, naively counting all pairs of replicates makes the benchmarking metrics depend on only a small number of compounds, which are principally positive controls. Balanced sampling metrics - where the number of pairs contributed a particular compound is capped at 100, which are sampled randomly - allows the benchmarking to depend on far more compounds. (A) The fraction of pairs (y-axis) rises very quickly with only a small fraction of compounds sorted by their number of pairs (x-axis). (B) Balanced sampling requires more compounds to achieve a particular pair fraction, meaning the benchmarking is a less biased assessment of assay performance. The CDRP dataset is relatively unaffected by balanced sampling, as only one compound had more than 100 pairs of signatures.

**Table S1:** For each cell line from S2, this table shows the AUC of the raw pair counts, the balanced AUC; the number of compounds needed to account for at least 50% of the pairs, the number of compounds with 50% of the pairs when using the balanced sampling, and the total number of compounds. The raw data requires as few as 2 compounds to account for half of the total pairs. Balanced sampling has lower AUC and requires more compounds to achieve any particular fraction of pairs. A dataset where each compound was assayed number of times would have an AUC of 0.5, and each compound would contribute equally to the benchmarks.

| cell_id | AUC | balAUC | N50 | balN50 | Total |
| --- | --- | --- | --- | --- | --- |
| A375 | 0.948 | 0.812 | 2 | 805 | 5048 |
| A549 | 0.971 | 0.851 | 2 | 833 | 6974 |
| ASC | 0.781 | 0.725 | 54 | 77 | 289 |
| BT20 | 0.693 | 0.632 | 54 | 65 | 186 |
| HCC515 | 0.913 | 0.847 | 40 | 259 | 3143 |
| HEK293 | 0.891 | 0.599 | 2 | 550 | 1388 |
| HEPG2 | 0.901 | 0.806 | 30 | 289 | 2523 |
| HUVEC | 0.824 | 0.677 | 27 | 234 | 818 |
| JURKAT | 0.684 | 0.651 | 412 | 466 | 1424 |
| MCF7 | 0.972 | 0.844 | 3 | 1015 | 7913 |
| NPC | 0.901 | 0.809 | 52 | 144 | 968 |
| PC3 | 0.973 | 0.867 | 2 | 898 | 8546 |
| VCAP | 0.923 | 0.753 | 6 | 887 | 14557 |
| CDRP | 0.628 | 0.628 | 4989 | 4991 | 15139 |

**Fig. S3:**
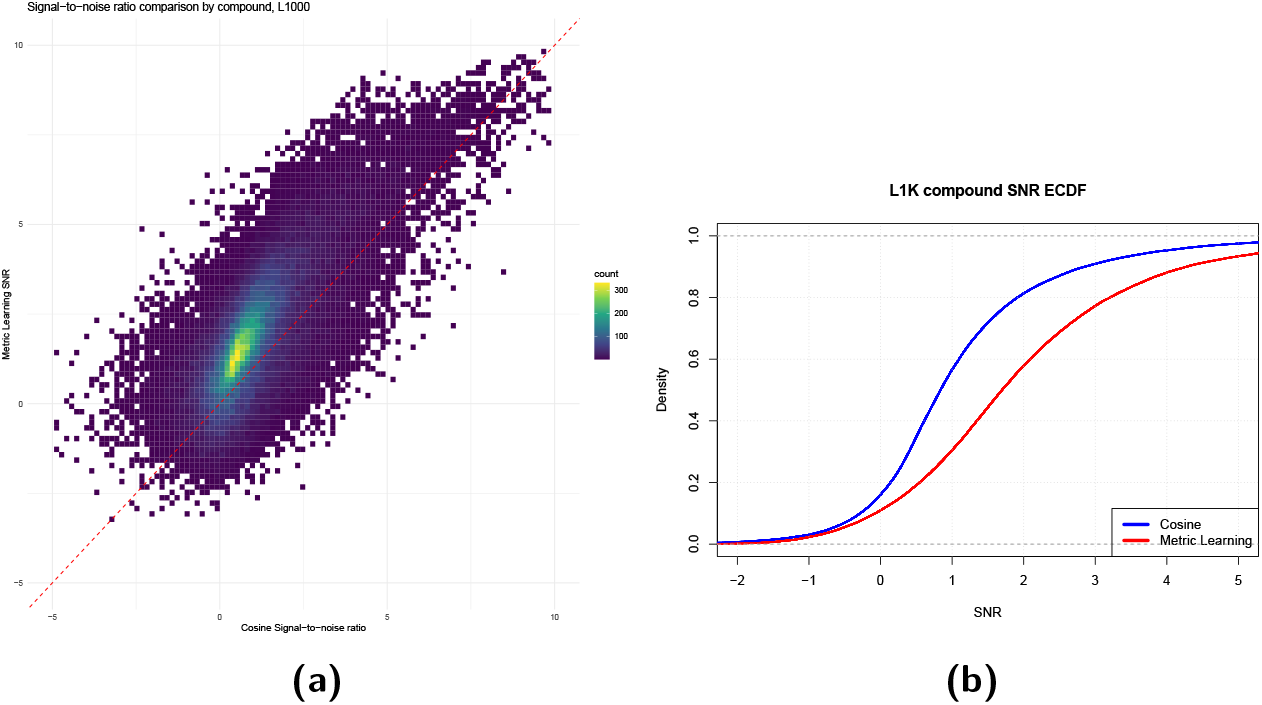
The signal-to-noise ratio of L1000 compound replicate similarities is greater with metric learning than with cosine.

**Fig. S4:**
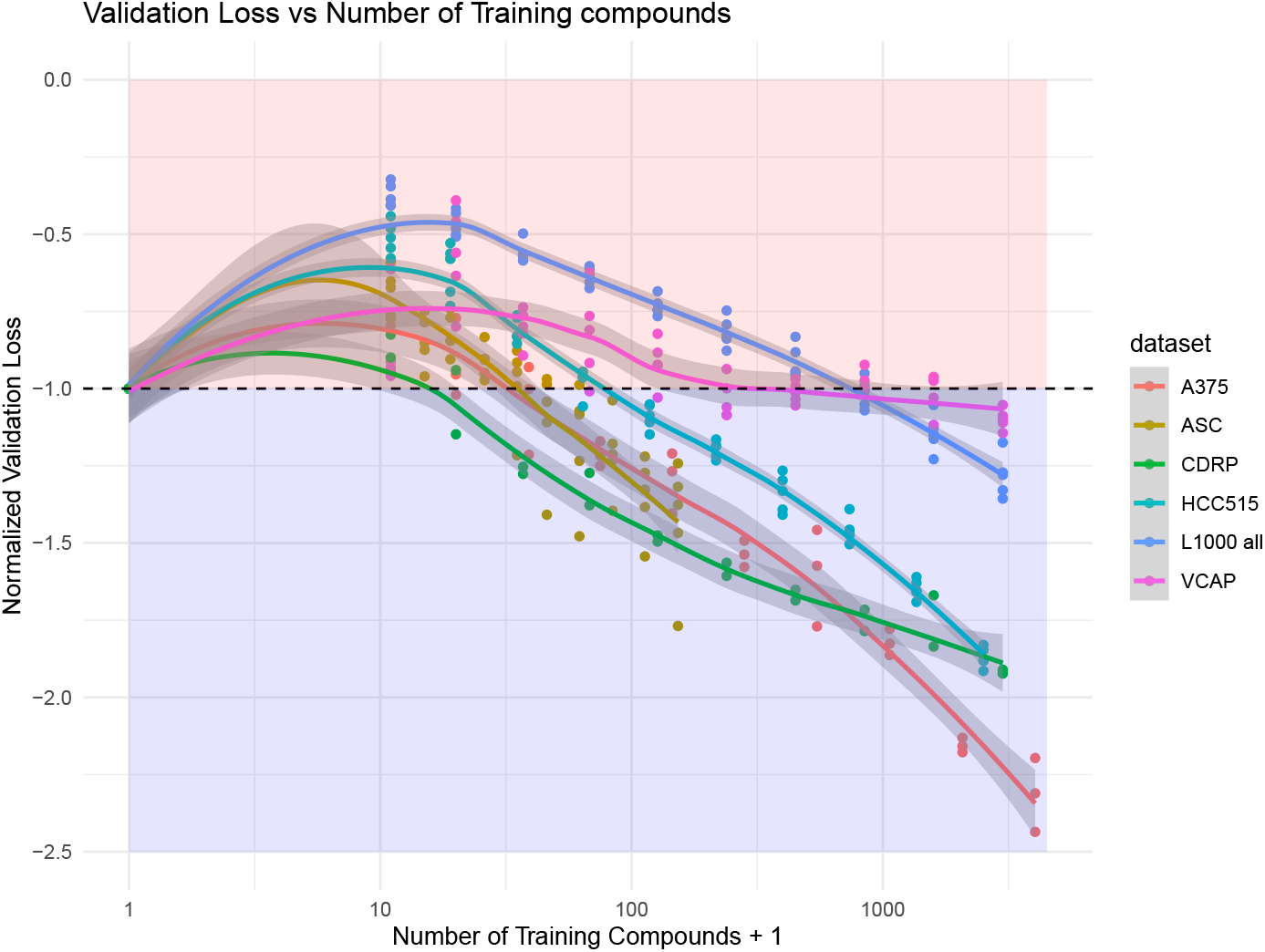
Validation t-loss as a function of the number of training compounds. A model is trained on N training compounds (x-axis), then the replicate loss is evaluated on held out test data, where no compounds in the test data are present in the training data. This process is repeated several times for each dataset or cell line, and the loss (y-axis) is evaluated on the test data. Smaller values of loss indicate better performance. The loss is scaled so a value of *−*1 corresponds to cosine similarity (equivalent to N=0 training examples). The blue region indicates regimes where metric learning outperforms cosine, and the red region where cosine outperforms metric learning. For all datasets in L1000 (cell lines or the entire dataset) and for the cell painting CDRP dataset, with sufficiently many training examples, metric learning outperforms cosine.

**Fig. S5:**
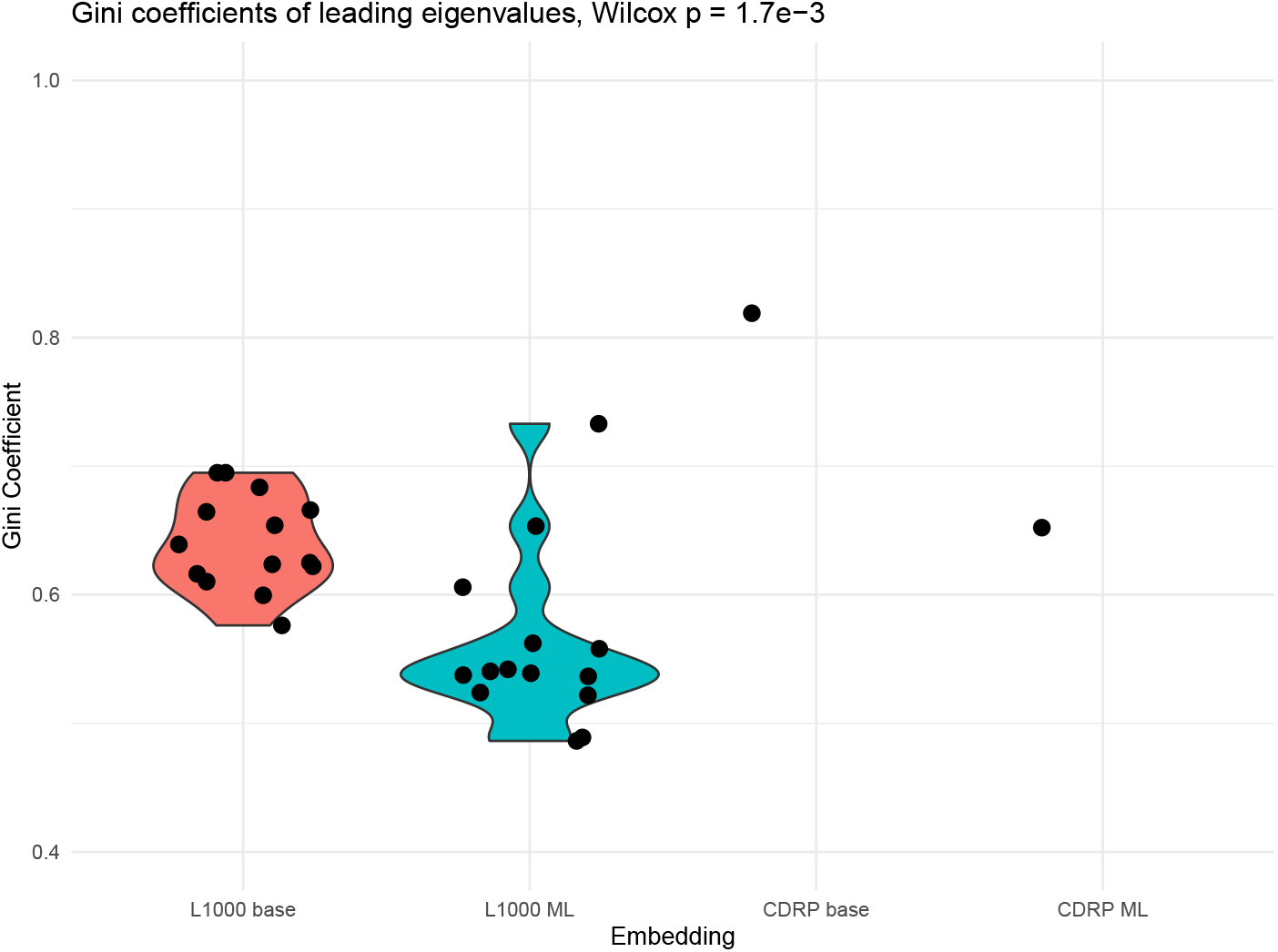
Gini coefficients of first 200 eigenvalues for L1000 and 100 eigenvalues for CDRP. The PeML embedding tends to lower the Gini coefficient of the eigenvalues, meaning that the first principal components contribute more equally to the variance. The gini coefficients of the ML embedding are statistically lower (p = 1.7*e −* 3 by paired Wilcoxon test)

